# Investigation of Antimicrobial and Antioxidant Activity of *Tephromela atra* Lichen and its Chemical Isolates, α-Alectoronic Acid and α-Collatolic Acid

**DOI:** 10.1101/2023.11.05.565661

**Authors:** Awad Alzebair, Meral Yilmaz Cankiliç, İlker Avan, Mehmet Candan

**Author notes:** Correspondence Author address: Awad ALZEBAIR^1^.

## Abstract

Lichens and their secondary metabolites are of particular interest as they can be used as an alternative to synthetic drugs in the treatment of various diseases. The present study was carried out to investigate in vitro antioxidant and antimicrobial activities of acetone, dichloromethane, petroleum ether, and methanol extracts of the lichen *Tephromela atra* and its major metabolites. α-alectoronic acid and α-collatolic acid were identified as bioactive substances in the lichen *Tephromela atra*. The antibacterial activity was investigated using the disk diffusion method and the minimum inhibitory concentrations were determined against eleven bacterial species, nine fungal species, and four yeast species using broth microdilution method. The antioxidant activity of the methanol extracts of *Tephromela atra* was measured by the free radical scavenging ability using the DPPH method. The extracts of α-alectoronic acid and α-collatolic acid showed significant inhibitory activity against most of the tested bacteria, especially the lowest MIC values of the two acids for *Bacillus subtilis*. Furthermore, the antioxidant activity was determined *in vitro* by measuring the radical scavenging capacity of the extracts on DPPH. The methanol extracts showed significant antioxidant activity and high DPPH scavenging activity at all concentrations. The results indicate that the extracts of *Tephromela atra* and the natural products obtained can be used as a source of antimicrobial and antioxidant agents in the pharmaceutical industry.

**HIGHLIGHTS:**

1. The present study determines the antibacterial and antifungal activities of the *Tephromela atra*, lichen extracts, and its isolated major constituents (α-alectoronic acid and α-collatolic acid), as well as evaluates the antioxidant capacity.
2. Although there are some studies on α-alectoronic acid and α-collatolic acid, isolated in different lichens, there is no study on the biological activity and antioxidant capacity of lichen *Tephromela atra*.
3. The MIC values of isolates were evaluated against a broad range of microorganisms including eleven species of bacteria, nine species of fungi, and four species of yeasts.
4. α-Alectoronic and α-collatolic acid extracts showed significant inhibitory activity against most of the tested bacteria, especially the lowest MIC values of both acids for Bacillus subtilis.
5. Our results provide a future perspective that *Tephromela atra* extracts and the isolated metabolites might be used as effective antimicrobial and antioxidant agents in pharmaceutical areas.

**Graphical Abstract:** 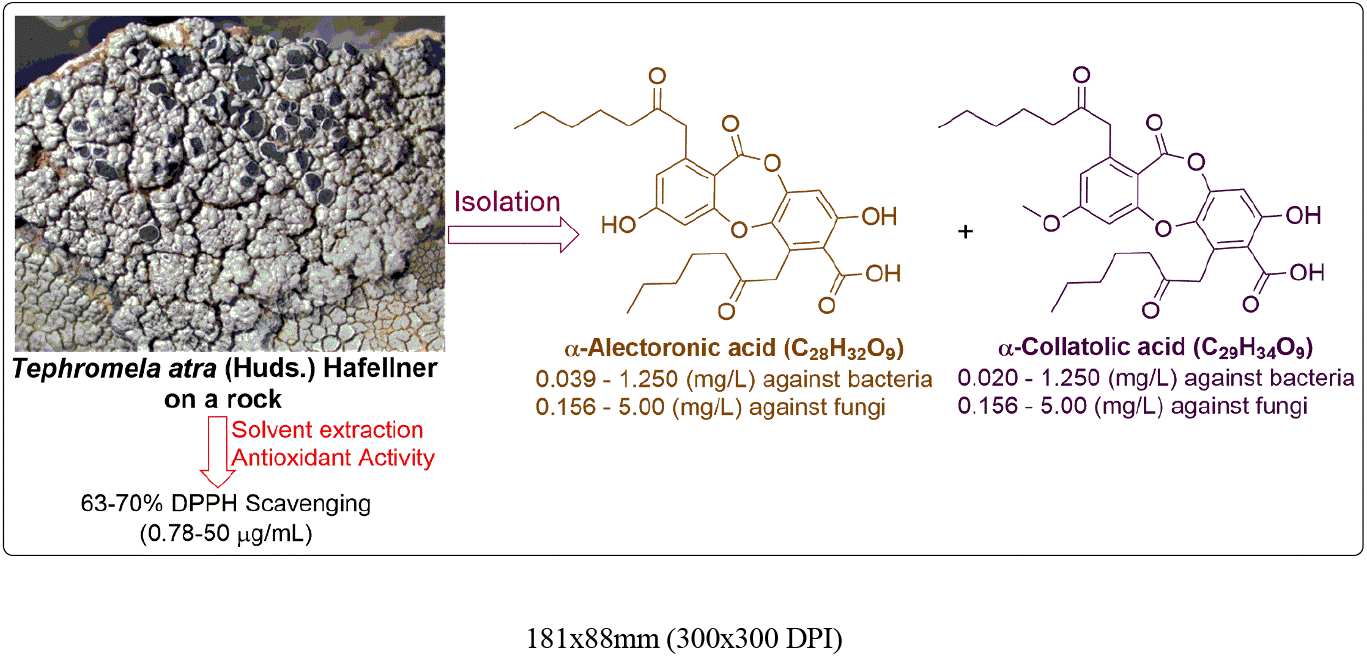

## INTRODUCTION

Scientists have long been interested in the use of new remedies of natural origin to treat diseases in plants, animals, and humans. It is known that the long-term use of synthetic drugs has some side effects and, above all, that the improper use of antibiotics makes microorganisms resistant to these drugs[1-3]. Lichens and their secondary metabolites are also an important group that can be used as an alternative to synthetic drugs in the treatment of various diseases[4]. Lichens are a unique group that evolved from a symbiotic relationship between photosynthetic organisms (cyanobacteria or algae) and fungi. There are about 20,000 taxa distributed around the world[5]. Lichens are used in industry and medicine for different purposes and in different ways. It is known that some lichen samples have long been used in the production of food, paint, alcohol, and perfume[6, 7]. The use of lichens in the treatment of various infectious diseases, especially in the field of traditional medicine, dates back to ancient times[8-12]. Many studies have shown that lichen metabolites have antimicrobial, antioxidant, anticarcinogenic, anti-inflammatory, antiproliferative, and cytotoxic effects[13-19].

The present study aims to evaluate the antioxidant capacity and determine the *in vitro* antibacterial and antifungal activities of the acetone, dichloromethane, petroleum ether, and methanol extracts of the lichen *Tephromela atra*, and its major constituents (α-alectoronic acid and α-collatolic acid). *Tephromela* genus has about 25-30 species and the most common species is *Tephromela atra* which has a worldwide distribution. Similarly, *Tephromela atra* has widespread dispersal areas in Turkey, and it contained many records from Eskisehir. It is a cosmopolitan species that develops on siliceous and less calcareous, nutrient-rich rocks and walls, rarely bark and timber. Although there are some studies on the listed compounds (α-alectoronic acid and α-collatolic acid), there is no study on the biological activity and phytochemical contents of *Tephromela atra*[20-23].

## MATERIAL AND METHODS

### Collection of lichen material

The lichen *Tephromela atra* (Huds.) Hafellner (Figure 1) was collected from an open area on the siliceous rock in Bozdag mountain in Eskisehir province in September 2018. and identified by lichenologist M. Candan, one of the authors of this study. Species identification was based on identification keys, flora books, and monographs. The lichen samples were deposited and stored in the herbarium of Biology Department of Eskisehir Technical University

**Figure 1.**
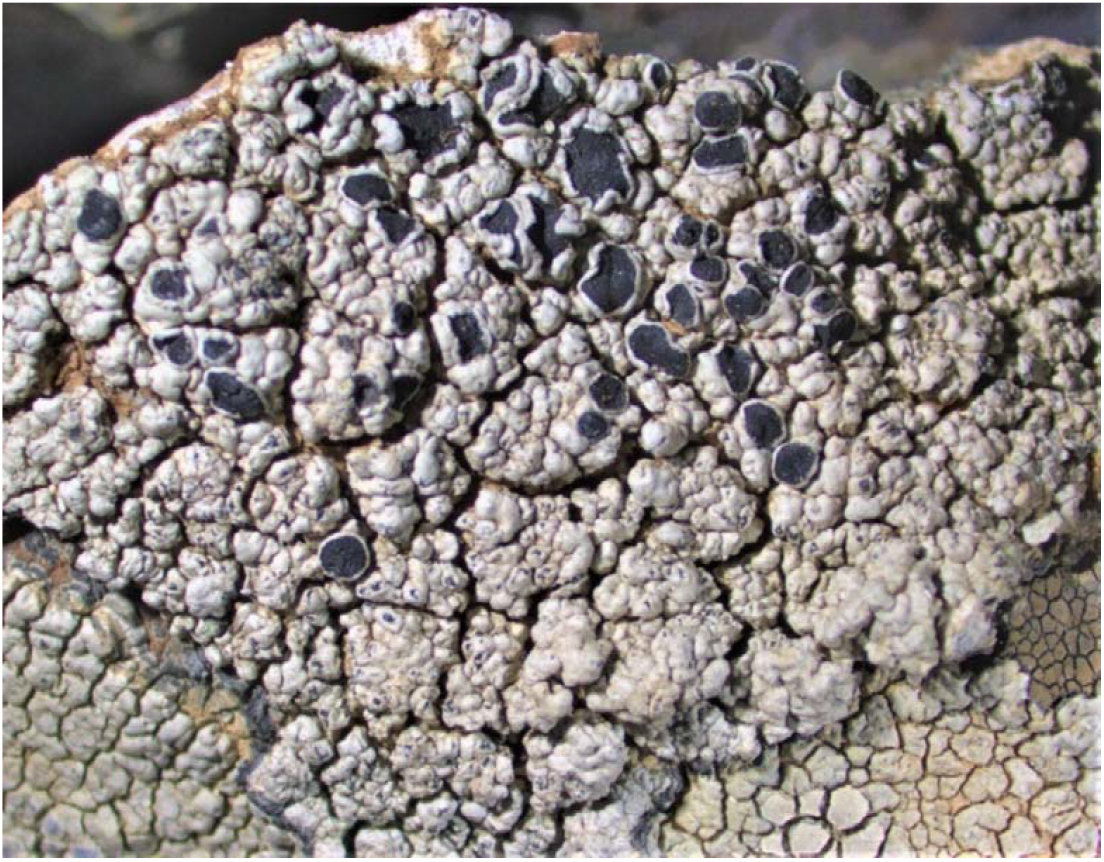
Microscopic image of *Tephromela atra* 357x278mm (118x118 DPI)

### Preparation of the lichen total extracts

A dry sample of *Tephromela atra* was first grounded, and 20 g portions of the sample was treated separately with solvents (100 mL) of different polarity, including petroleum ether, dichloromethane, acetone, and methanol. The mixtures were first sonicated in ultrasound apparatus at 38 °C for 20 minutes and then left overnight at room temperature for 24 hours. After the mixtures were filtered off, the filtrates were concentrated and dried under reduced pressure to obtain dry total extracts. The dry extracts were kept at 4 °C.

### Microorganisms and media

The antimicrobial activity of lichen extract was studied on the following bacteria, yeasts, and filamentous fungi: *Listeria monocytogenes* ATCC 19111, *Bacillus cereus* ATCC 10876, *Bacillus subtilis* NRRL NRS-744, *Micrococcus luteus* NRRL B-4375, *Enterobacter aerogenes* NRRL 3567, *Streptococcus faecalis* NRRL B-14617, *Escherichia coli* ATCC 25922, *Proteus vulgaris* NRRL B-123, *Klebsiella pneumoniae* ATCC 700603, *Yersinia enterocolitica* Y53, and *Staphylococcus aureus* ATCC 6538 bacteria. *Penicillium notatum, Penicillium expansum, Penicillium citrinum, Fusarium solani* ATCC 12820, *Fusarium moniliforme* NRRL 2374, *Aspergillus fumigatus* NRRL 113, *Aspergillus flavus* ATCC 9807, *Aspergillus niger* ATCC 6275, *Aspergillus ochraceus* filamentous fungi. *Candida albicans* ATCC 90028, *Candida krusei* ATCC 6258, *Candida globrata* ATCC 90030, *Candida parapsilosis* ATCC 22019 yeasts. The microorganisms were obtained from Microbiology Research Laboratory, Department of Biology, Faculty of Science, Eskisehir Technical University. Cultures of each of the bacteria were maintained on Mueller Hinton Agar (MHA) as it is a standard medium best suited for susceptibility testing using the Kirby-Bauer disc diffusion method[24, 25], and the cultures of fungi were maintained on Sabouraud Dextrose Agar (SDA).

### In vitro antimicrobial activity

The total extracts obtained from the above solvents were again added to 100 mL of the previous solvents and the mixtures were homogenized using a vortex mixer and sonicator. Then 50 blank sterile antibiotic discs (7 mm diameter) were soaked in 1 mL of these solutions and homogenized by shaking. The solvents on the antibiotic discs were evaporated in a vacuum oven while shaking. The amount of total extracts, concentrations of solutions, and the amount of absorbed total extract on each disc were listed in Table 1.

**Table 1.**
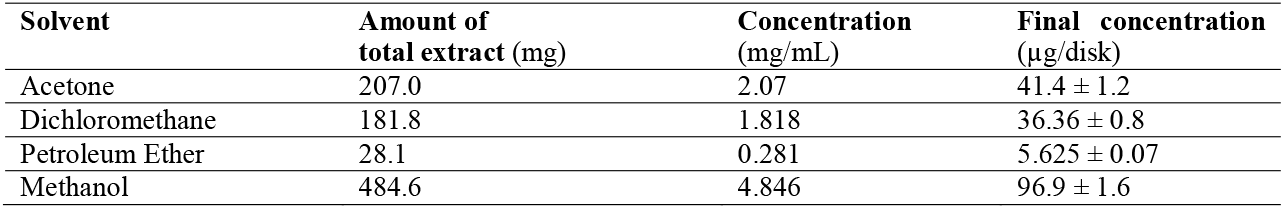
The amount of total extracts (mg), concentrations of solutions (mg/mL), and the amount of absorbed total extract on each disc (μg/disk). Mean value of 3 experiments ± standard deviation (SD).

The anti-microbial activity of extracts was determined by using the disk diffusion method. For the disk diffusion method, antimicrobial susceptibility was tested on MHA solid media in Petri dishes. The bacteria from tested microorganisms were activated by incubation in 10 mL Mueller-Hinton Broth (MHB) medium at 30-37° C for 24–48 hours. Yeasts were activated by incubation in SD Broth medium at 37°C for 24–48 hours. The concentration of activated bacteria and yeasts was adjusted according to McFarland No: 0.5 (10^8^ CFU/ml). Filamentous fungi were developed on potato dextrose (PD) agar for 10 days at 25°C to obtain their spores, and 5 mL of sterile distilled water containing 0.1% Tween 80 was poured onto the developing molds. After waiting for 20-30 minutes, the spore suspension obtained was counted on the Thoma slide and the number of spores was determined and adjusted to 10^5^ spores/mL.

The sensitivity of the microorganisms to the different concentrations of the acetone, dichloromethane, methanol, and petroleum ether extracts was tested using the disk diffusion method by Kirby and Bauer by measuring the zone of inhibition of the antibiotic disks (7 mm diameter) containing the extracted substance[24, 25]. The blank disc was used as a negative control on the plates. MH, SD and PD agar were seeded with the appropriate inoculum for bacteria, yeasts, and filamentous fungi, respectively. Paper disks (7 mm) containing lichen extracts were placed on the test plates using sterile forceps. The agar plates were cultivated at 37 °C for 24–48 hours for bacteria and at 25 °C for 5–7 days for filamentous fungi. Chloramphenicol and ketoconazole were used as controls for bacteria and fungi, respectively. Finally, the antimicrobial activities of the different extracts were determined by observing the inhibitory zones around the disks[26, 27]. All experiments were performed in triplicate.

### Isolation of α-alectoronic acid, and α-collatolic acid

The lichen (*Tephromela atra*) sample was first crushed, and then a 20 g portion of the dry sample was soaked in 250 mL acetone. The mixture was placed in a sonicater for 30 minutes and then kept overnight at room temperature in the dark. The mixture was filtered through a 10 cm silica pad to remove physical dirt. The existing lichen acids present in the extract of *Tephromela atra* were identified by comparing their Rf values with reported values in A, C, and G eluent systems[9, 28, 29]. Most of the solvent was evaporated under reduced pressure using a rotary evaporator. The lichen acids (Figure 2), α-alectoronic acid, and α-collatolic acid were isolated from the crude extract solution using preparative thin-layer chromatography (TLC) and a modified solvent G mix (toluene/ethyl acetate/formic acid (139: 83: 8 v/v/v)), as similarly described for the lichen, Hypogymnia physodes (L.) Nyl[30]. In a typical TLC separation, 50–100 mg of *Tephromela* crude extract was dissolved in acetone (10 mL) to prepare a stock solution. 2–3 mL of stock solution was applied to preparative TLC plates (glass, Merck, 20x20 cm, and 0.25 mm thick silica-60 coated with F254, a fluorescent indicator). The crude-loaded TLC plates were developed twice using a modified solvent G system. The development and separation were monitored under UV light (254 nm). After satisfactory development, the TLC plates were removed from the chamber. After the plates were dried, the lichen acid bands were scraped off separately from the plates. The acid adsorbed silicas were extracted with acetone (2 × 50 mL). The silica portion was removed by filtration. The solvents were evaporated. The residues were recrystallized over acetone to obtain pure products. The TLC separations were repeated until a sufficient amount of compounds was obtained for the antimicrobial tests. Characterization of α-alectoronic acid and α-collatolic acid was based on high-resolution mass spectroscopy (HRMS). The HRMS analyses were performed using a Shimadzu LC-MS IT-TOF HRMS system. HRMS [ESI(+)-TOF] of α-alectoronic acid: m/z [M + H]+ calcd. for C28H32O9: 513.2119; found: 513.2117. HRMS [ESI(+)-TOF] of α-collatolic acid: m/z [M + H]+ calcd. for C29H34O9: 527.2276; found: 527.2273 (Supplementary Information).

**Figure 2.**
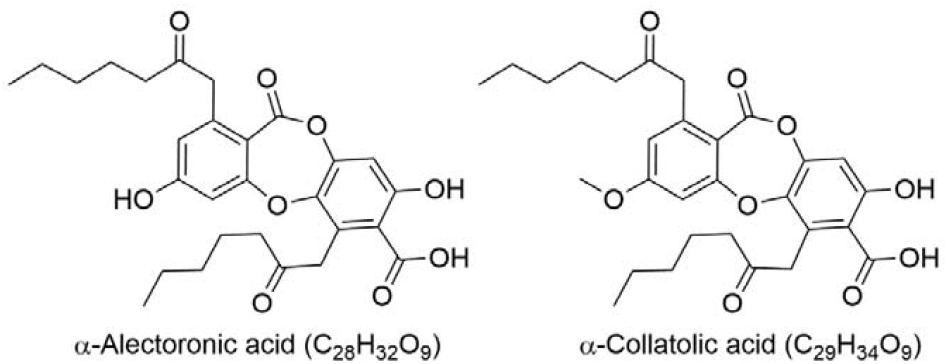
Chemical structures of α-alectoronic acid and α-collatolic acid. 120x46mm (300x300 DPI)

### Minimal inhibitory concentration (MIC) of α-alectoronic acid and α-collatolic acids

The minimum inhibitory concentrations of the isolated lichen acids, α-alectoronic acids and α-collatolic acids, were performed using the broth microdilution method with 96-well microtiter plates. The concentrations of α-alectoronic acid and α-collatolic acid in the medium were adjusted in the range of 10 mg/mL and 19.5 μg/mL, respectively. Before the dilution process, the lichen acids were dissolved in DMSO (20% in water) and their starting solutions were obtained first. The ten diluted test tubes were prepared with a two-fold serial dilution technique using MHB for bacterial cultures, SD broth, and PD broth. Then, 100 μl of each dilution plus 100 μL activated microorganism solution at 10^8^ cells/mL or fungal spore suspensions at 10^5^ spores/mL were transferred to microtitration plates and incubated for 24–48 h at 30–37 °C for bacteria/yeasts and at 28 °C for one week with daily observation for filamentous fungi. While the wells containing medium and microorganisms were used as a positive control group, the wells containing only medium served as a negative control. A third group was set up as a control with ketoconazole for fungi and chloramphenicol for bacteria. The positive and negative results were evaluated by the turbidity that occurred after 24–48 hours compared to the control wells. The lowest concentrations providing no observable growth of each microorganism were assigned as MIC. A 0.5% TTC (2,3,5-triphenyltetrazolium chloride, Merck) aqueous solution was applied to determine the MIC values to indicate the lowest concentration doses at which the microorganisms grew as a limiting dilution without color change[31, 32].

### Antioxidant activity by scavenging DPPH radicals

The antioxidant activity of the methanol extracts of *Tephromela atra* was measured based on the ability to capture the free radical by using 1,1-diphenyl-2-picryl-hydrazyl (DPPH)[16]. The analysis was performed according to the reported method described by Yu et al[33]. First, 0.6 mM of DPPH in methanol solution was prepared by dissolving of DPPH (0.024 g) into 100 mL of methanol. Then, 10 mL of this solution was diluted to 100 mL with methanol to prepare the final stock solution (0.06 mM) of DPPH, which was stored in dark at room temperature. Second, *Tephromela atra* extracts at varying concentrations (10, 20, 40, 60, 80, 100, 120, 140 μg/mL) were tested and ascorbic acid was used as a reference standard for the determination of the antioxidant activity by DPPH method[34]. The concentration of ascorbic acid varied from 1 to 60 μg/mL. A volume of 100 μL of each different concentration of *Tephromela atra* methanolic extract sample was mixed with 200 μL of 0.06 mM methanolic solution of DPPH in a 96-well plate, and the plate was incubated at room temperature in dark for 30 minutes, after that, the absorbance was measured at 517 nm on the spectrophotometer[35]. The measurements were repeated three times for each extract and the results were averaged. The free radical scavenger of the extract solutions is indicated as % inhibition of DPPH absorption. IC_50_ was calculated from % inhibition. The calculation of the remaining DPPH was done as given in the following equation.

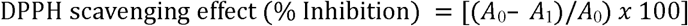

A_0_ = The absorption of the control sample.

A_1_ = The absorption in the existence of all extract samples or references.

Statistical calculations were performed using the EXCEL and simple linear regression analysis was utilized to determine the statistical significance of extract antioxidant activity.

## RESULTS

The study includes the screening of antibacterial and antifungal activity of acetone, dichloromethane, petroleum ether, and methanol extracts of *Tephromela atra* lichen against eleven Gram-positive and Gram-negative bacteria, nine filamentous fungi, and four yeasts using the disk diffusion method (Table 2 and Table 3). The results in Table 2 show that the extracts of *Tephromela atra* have antibacterial activity against most of the tested microorganisms. The acetone and dichloromethane extracts are effective against all tested bacteria except *K. pneumoniae*, while the petroleum ether extract showed activity against all tested bacteria except *E. aerogenes, S. faecalis, K. pneumoniae*, and *Y. enterocolitica*. The methanol extract showed activity against all bacteria tested with the exception of *E. aerogenes* and *Y. enterocolitica*. With the exception of the petroleum ether extract, the other extracts showed antibacterial activity against *S. faecalis* bacteria.

**Table 2.** Antibacterial activities of *Tephromela atra* extracts against different Gram-positive and Gram-negative bacteria according to the disk diffusion test. (+): show inhibitory against tested microorganism (-): show no inhibitory against tested microorganism. Mean value of 3 experiments ± standard deviation (SD).

| **Bacteria** | **Acetone**<br>(41.4 ± 1.2 μg/disk) | **Dichloromethane**<br>(36.36 ± 0.8 μg/disk) | **Petroleum Ether**<br>(5.625 ± 0.07 μg/disk) | **Methanol**<br>(96.9 ± 1.6 μg/disk) |
| --- | --- | --- | --- | --- |
| *E. aerogenes* | + | + | - | - |
| *E. coli* | + | + | + | + |
| *P. vulgaris* | + | + | + | + |
| *K. pneumoniae* | - | - | - | + |
| *Y. enterocolitica* | + | + | - | - |
| *B. cereus* | + | + | + | + |
| *B. subtilis* | + | + | + | + |
| *L. monocytogenes* | + | + | + | + |
| *M. luteus* | + | + | + | + |
| *S. faecalis* | + | + | - | + |
| *S. aureus* | + | + | + | + |

**Table 3.** Antifungal activities of *Tephromela atra* extracts against different filamentous fungi and yeasts according to the disk diffusion test. (+): show inhibitory against tested microorganism (-): show no inhibitory against tested microorganism. Mean value of 3 experiments ± standard deviation (SD).

| **Microorganisms** | **Acetone**<br>(41.4 ± 1.2<br>μg/disk) | **Dichloromethane**<br>(36.36 ± 0.8<br>μg/disk) | **Petroleum Ether**<br>(5.625 ± 0.07<br>μg/disk) | **Methanol**<br>(96.9 ± 1.6<br>μg/disk) |
| --- | --- | --- | --- | --- |
| **Yeasts** |  |  |  |  |
| *C. globrata* | - | - | - | - |
| *C. parapsilosis* | - | - | - | - |
| *C. krusei* | + | + | + | + |
| *C. albicans* | - | - | - | - |
| **Filamentous fungi** |  |  |  |  |
| *A. flavus* | - | - | + | - |
| *A. fumigatus* | - | - | + | + |
| *A. ochraceus* | + | - | + | + |
| *A. niger* | - | - | + | + |
| *F. moniliforme* | - | - | + | - |
| *F. solani* | - | - | + | + |
| *P. notatum* | + | - | + | - |
| *P. expansum* | - | - | + | - |
| *P. citrinum* | + | - | + | - |

The results in Table 3 show that all extracts have anti-yeast activity against *Candida krusei*. The petroleum ether extract showed antifungal activity against all tested filamentous fungi. The acetone extract, on the other hand, is effective against *A. ochraceus, P. notatum* and *P. citrinum*, while the methanol extract is effective against *A. fumigatus, A. ochraceus, A. niger*, and *F. solani*. It can be clearly seen that the dichloromethane extract is only effective against bacteria rather than fungi, with the exception of *C. krusei* yeast.

The minimum inhibitory concentration values of α-alectoronic acid and α-collatolic acid (Table 4) were determined against the same test microorganisms using the broth microdilution method with 96-well microtiter plates[25]. The results show that α-alectoronic acid and α-collatolic acid from *Tephromela atra* have a significant effect against most microorganisms at different concentrations.

**Table 4.** MIC values (μg/mL) of α-alectoronic acid and α-collatolic acid of *Tephromela atra* against bacteria. Mean value of 3 experiments ± standard deviation (SD).

| **Microorganisms** | **α-Collatolic acid** | **α-Alectoronic acid** | **Chloramphenicol** |
| --- | --- | --- | --- |
| **Bacteria** | **MIC ($\mu\text{g/mL}$)** | **MIC ($\mu\text{g/mL}$)** | **MIC ($\mu\text{g/mL}$)** |
| *E. aerogenes* | $312.5 \pm 2.1$ | $39 \pm 0.3$ | $78.1 \pm 0.6$ |
| *E. coli* | - | - | $19.5 \pm 0.2$ |
| *P. vulgaris* | $156.2 \pm 1$ | $78.1 \pm 0.6$ | $39 \pm 0.3$ |
| *K. pneumoniae* | - | - | $78.1 \pm 0.6$ |
| *Y. enterocolitica* | $39 \pm 0.3$ | $78.1 \pm 0.6$ | $78.1 \pm 0.6$ |
| *B. cereus* | $39 \pm 0.3$ | $78.1 \pm 0.6$ | $39 \pm 0.3$ |
| *B. subtilis* | $39 \pm 0.3$ | $19.5 \pm 0.2$ | $19.5 \pm 0.2$ |
| *L. monocytogenes* | - | - | $39 \pm 0.3$ |
| *M. luteus* | $78.1 \pm 0.6$ | $39 \pm 0.3$ | $19.5 \pm 0.2$ |
| *S. faecalis* | $1250 \pm 4.2$ | $312.5 \pm 2.1$ | $39 \pm 0.3$ |
| *S. aureus* | $78.1 \pm 0.6$ | $78.1 \pm 0.6$ | $78.1 \pm 0.6$ |

α-Collatolic acid showed equipotent activity to the standard antibiotic chloramphenicol against *B. cereus* (MIC= 39 μg/mL) and *S. aureus* (MIC= 78.12 μg/mL). It also showed antibacterial activity against *B. subtilis* with half the potency of chloramphenicol (MIC= 39 μg/mL). Compared to chloramphenicol, the highest antibacterial activity of α-collatolic acid was obtained against *Y. enterocolitica* (MIC= 39 μg/mL), while the lowest antibacterial activity was recorded against *S. faecalis* (MIC= 1250 μg/mL).

α-alectoronic acid had equipotent activity to the standard antibiotic chloramphenicol against *Y. enterocolitica, S. aureus* (MIC= 78.12 μg/mL) and *B. subtilis* (MIC= 19.5 μg/mL). The MIC value obtained against *E. areogenes* (MIC= 39 μg/mL) was lower than that of chloramphenicol. However, it showed moderate antibacterial activity against *B. cereus, P. vulgaris* (MIC= 78.12 μg/mL), and *M. luteus* (MIC= 39 μg/mL) with half the potency of chloramphenicol. The lowest antibacterial activity of α-alectoronic acid was against *S. faecalis* (MIC= 312.5 μg/mL) similar to α-collatolic acid. As can be seen in Table 4, no correlation was found between the substances and the activities of α-collatolic acid and α-alectoronic acid. Table 5 contains the results of the MIC values for α-alectoronic acid and α-collatolic acid extracts against filamentous fungi and yeasts.

**Table 5.** MIC values (μg/mL) of α-alectoronic acid and α-collatolic acid of *Tephromela atra* against fungi and yeasts. Mean value of 3 experiments ± standard deviation (SD).

| Microorganisms | α-alectoronic acid | α-collatolic acid | Ketoconazole |
| --- | --- | --- | --- |
| Fungi | MIC ($\mu\text{g/mL}$) | MIC ($\mu\text{g/mL}$) | MIC ($\mu\text{g/mL}$) |
| *F. moniliforme* | - | 2500 ± 17.1 | 39 ± 0.3 |
| *F. solani* | - | - | 78.12 ± 0.6 |
| *P. citrinum* | 2500 ± 17.1 | 156 ± 1 | 78.12 ± 0.6 |
| *P. expansum* | - | - | 78.12 ± 0.6 |
| *P. notatum* | 2500 ± 17.1 | 625 ± 7.3 | 39 ± 0.3 |
| *A. niger* | 2500 ± 17.1 | 2500 ± 17.1 | 78.12 ± 0.6 |
| *A. flavus* | 1250 ± 4.2 | 5000 | 78.12 ± 0.6 |
| *A. ochraceus* | 2500 ± 17.1 | 625 ± 7.3 | 78.12 ± 0.6 |
| *A. fumigatus* | - | - | 78.12 ± 0.6 |
| **Yeasts** |  |  |  |
| *C. albicans* | 2500 ± 17.1 | 156 ± 1 | 78.12 ± 0.6 |
| *C. krusei* | 5000 | 625 ± 7.3 | 78.12 ± 0.6 |
| *C. glabrata* | - | 1000 ± 3.9 | 39 ± 0.3 |
| *C. paropilopsis* | 156 ± 1 | 156 ± 1 | 78.12 ± 0.6 |

When evaluating the results of the antifungal activity, it became clear that the MIC values achieved were quite high compared to the standard antibiotic ketoconazole. α-collatolic acid had moderate antifungal activity against filamentous fungi *P. citrinum* (MIC=156 μg/mL) and *C. albicans, C. paropilopsis* (MIC= 156 μg/mL) yeasts with half the potency of ketoconazole. α-alectoronic acid also showed moderate antifungal activity against *C. paropilopsis* (MIC= 156 μg/mL) at half potency of ketoconazole. From Tables 4 and 5, it was understood that the antimicrobial effects of these lichen acids were higher against bacteria than fungi.

The antioxidant activity of methanol extracts was measured applying 1,1-diphenyl-2-picrylhydrazyl (DPPH) method, and the free radical scavenger of the extract solutions was indicated as % inhibition of DPPH absorption. The DPPH scavenging effect (% inhibition) calculated according to equation 1 and then IC50 values of the extract (35.23 μg/mL) and ascorbic acid (7,12 μg/mL) were determined using the regression equation. Ascorbic acid at a concentration of 10 μg/mL exhibited a percentage inhibition of 55.78% and for 60 μg/mL 99.68% [Table 6]. Figure 3 shows how the reducing power of the test extracts increases with the increase in amount of sample. The reducing power shows a good linear relationship in both ascorbic acid (R^2^ =0,9409) and MeOH extract (R^2^= 0.9602). It was observed that methanolic extracts show significant DPPH radical scavenging property. In addition, when the significance of the F statistic (F: 144,7947357; Significance F: 1,99864E-05) in the ANOVA table obtained by regression analysis was examined, it was concluded that the model had a good fit measure.

**Table 6.** DPPH scavenging 100% of different concentrations (ug/mL) of *Tephromela atra* methanolic extract and ascorbic acid.

| **Concentration**<br>**μg/mL** | **% inhibition of**<br>**methanol extract** | **% inhibition of**<br>**ascorbic acid** |
| --- | --- | --- |
| 10 | 35.42 | 55.78 |
| 20 | 48.6 | 68.86 |
| 40 | 55.3 | 86.88 |
| 60 | 58.7 | 99.68 |
| 80 | 65.6 |  |
| 100 | 71.9 |  |
| 120 | 76.1 |  |
| 140 | 88.2 |  |

**Figure 3.**
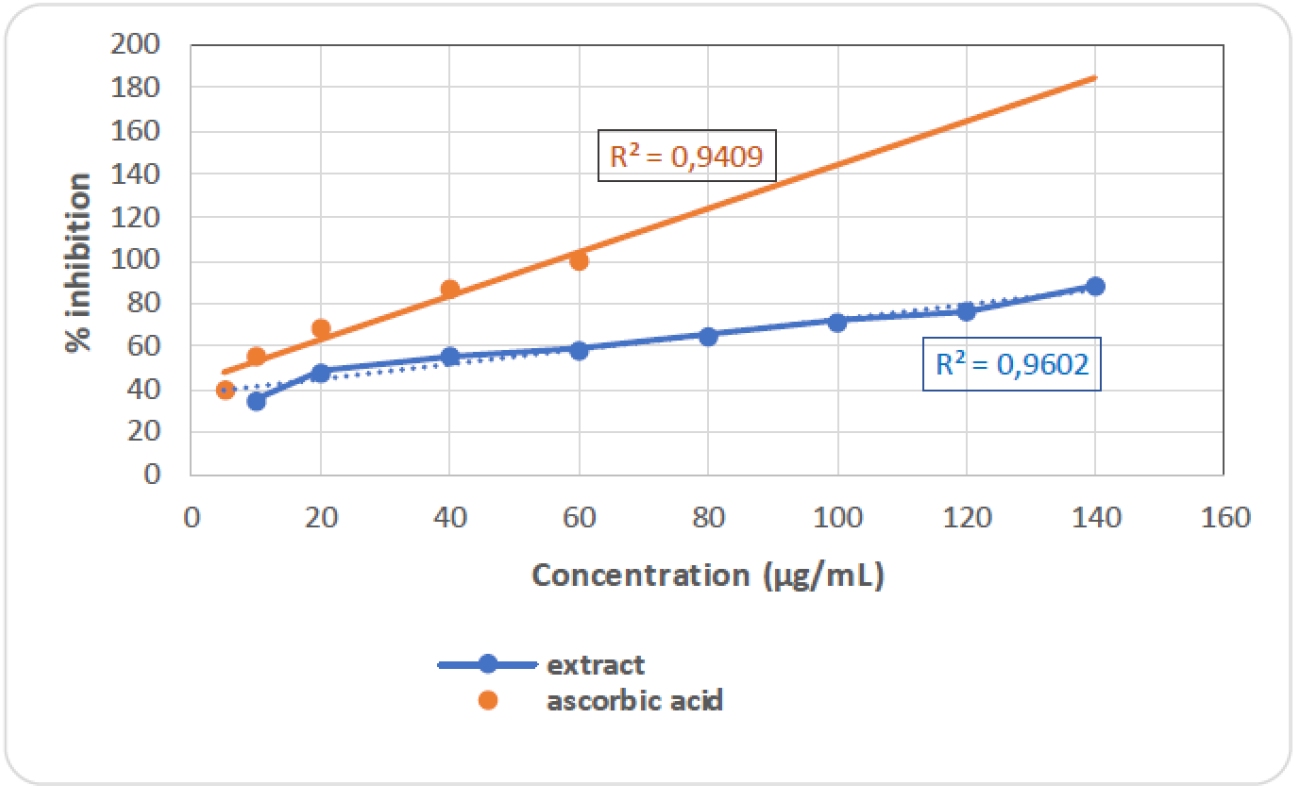
DPPH scavenging 100% to different concentrations (μg/mL) of *Tephromela atra* methanolic extract. 458x294mm (118x118 DPI)

## DISCUSSION

Increasing the number of antibiotic-resistant microorganisms due to unconscious use reduces the effectiveness of existing antibiotics. Further screening and investigation of natural pharmaceutical raw materials will help to reduce the problems of drug resistance. In contrast to plant species, lichens, as symbiotic organisms, synthesize a large number of chemical metabolites in large quantities. Lichens have also gained special importance because the acids and other metabolites of lichens are mostly unique and not found in other plant groups[25]. Therefore, recent studies have focused on lichen extracts, their metabolites and their pharmacological uses. As a result, many lichen metabolites have been characterized and their biological activities investigated. It is known that some of these metabolites are intrinsic antibiotics that protect the lichen organisms from micropredators[21, 36-38]. Enormous research has been conducted in many countries on the antibiotic properties of lichen metabolites. Today, many lichen substances have been identified as antibiotics. For example, atranorin, gyrophoric acid, fumarprotocetraric acid, lecanoric acid, physodic acid, protocetraric acid, stictic acid, usnic acid, etc. have been found to have relatively strong antimicrobial effects against various bacteria and fungi as plant, animal and human pathogens[2, 38-40].

In this study, acetone, dichloromethane, petroleum ether and methanol extracts obtained from the lichen *Tephromela atra* were tested for their antibacterial and antifungal activity against a variety of microorganisms using the disk diffusion method. In addition, α-alectoronic acid and α-collatolic acid were isolated from the same lichen using the TLC plate technique. The minimum inhibitory concentration (MIC) of the isolated metabolites against various microorganisms was determined by broth microdilution method with 96-well microtiter plates. The antioxidant activity of the methanolic extracts of the same lichen was analyzed by using the DPPH free radical method.

All extracts have shown activity against *E. coli, P. vulgaris, B. cereus, B. subtilis, L. monocytogenes, M. luteus, S. aureus*. Only the methanolic extract is effective against *K. pneumonia* and the acetone and dichloromethane extracts are effective against *E. aerogenes* and *Y. enterocolitica*. On the other hand, the results show that the petroleum ether extract of *Tephromela atra* has greater antifungal activity against most of the tested fungi than the extracts of acetone, methanol and dichloromethane. The dichloromethane extract of *Tephromela atra* showed no significant effect against most of the yeasts and filamentous fungi tested in this study, except against *C. krusei* ATCC 6258.

*S. aureus* is at the top of the list of dangerous pathogens responsible for nosocomial and community-acquired infections in the last 30 years[41, 42]. Vancomycin is the only option for the treatment of S. aureus infections, and this is the main reason for the search for new alternatives. In our study, *S aureus* ATCC 6538 type was also used as a test microorganism. It is worth mentioning that all the different extracts of *Tephromela atra* tested in our study showed antagonistic activity on *S. aureus*, which is clinically important, as shown in Tables 3 and 4. The extracts of *Tephromela atra*, α-alectoronic acid and α-collatolic are tested against a broad spectrum of microorganisms and their minimum inhibition concentrations are determined. The results show that α-alectoronic acid and α-collatolic acid from *Tephromela atra* have a significant effect against all tested microorganisms at different concentrations. Considering the remarkable information of this study, it is found that *Tephromela atra* extracts have impressive antibacterial activity against Gram-positive bacteria and inhibited the growth of Gram-negative bacteria. In addition, all extracts have remarkable anti-yeast activity against *C. krusei*. The MIC values of α-alectoronic acid and α-collatolic acid are also significant against the tested microorganisms at different concentrations, besides their free radical scavenging effect and antioxidant activity.

The biological activities of α-alectoronic acid and α-collatolic acid isolated from *Parmotrema* species were recently reported by Rajan et al[21]. In their study, α-alectoronic acid and α-collatolic acid had antibacterial activity against *S. aureus* (MIC value 500 μg/mL) and *B. subtilis* (MIC value 125 μg/mL). The MIC values we obtained were lower against S. aureus bacteria than in this study. Therefore, we extended our study to include a wide range of fungi and yeasts in addition to bacteria.

In a study by Piovano et al[18], the MIC of α-collatolic acid was found to be above 250 μg/ml against *Candida albicans, Saccharomyces cerevisae, Cryptococcus neoformans, Aspergillus flavus, A. niger, Microsporum canis* C 112, *Microsporum gypseum* C 115, *Trichophyton mentagrophytes* ATCC 9972, *T*.*rubrum* C 113 and 12.5 μg/mL for *Staphylococcus aureus* (methicillin sensitive), *Staphylococcus aureus* (methicillin resistant). The results of this study support the results of our study.

In another study by Verma et al.[20], some lichen compounds, including α-collatolic acid and α-alectoronic acid, isolated from different lichens (*C. ochrochlora, P. nilgherrensis* and *P. sanctiangelii*) showed moderate to strong bactericidal activity with low MIC values; α-alectoronic acid showed an MIC of 21.9 μg to 162.1 μg/ml and α-collatolic acid an MIC of 8.6–245 μg/ml against examined species. These values also supported our study.

The production of harmful free radicals, which overwhelm the body’s antioxidant defense capacity, leads to an increase in oxidative stress. The production of these reactive oxygen species is due in part to environmental factors such as pollution, temperature, excessive light intensity, and dietary restrictions[43]. These highly reactive free radicals are known to be responsible for some human diseases such as cancer and cardiovascular disease[44]. Therefore, there is a need for an external source of antioxidants to combat the effects of reactive free radicals, and lichens have been shown to possess promising antioxidant activity[45].

Numerous researchers have previously investigated the antioxidant activity in lichen extracts and metabolites[36, 46-48]. Using 1,1-diphenyl-2-picrylhydrazyl (DPPH) radical scavenging, reducing power, and superoxide anion free radical scavenging experiments, Kosanić et al.[14] showed that the acetone extracts of *Evernia prunastri* and *Pseudoevernia furfuraceae* lichens and their main metabolites (evernic acid and physodic acid) had strong antioxidant activity *in vitro*.

This is the first study to investigate the antioxidant activity of *Tephromela atra* extracts, and the results showed that methanol extracts of *Tephromela atra* highly scavenge DPPH free radicals. Therefore, we can conclude that methanol extracts of *Tephromela atra* are a promising source of antioxidants. In addition, the data from this study indicate that the extracts could be a potential precursor for the research and development of active substances against infectious diseases and free radical-induced oxidative stress. In this sense, it is quite important to continue and support studies to search for new compounds from lichens exhibiting strong antioxidant activity.

## CONCLUSION

The antimicrobial effects of these important acids isolated from *Tephromela atra* (α-alectoronic acid and α-collatolic acid) show strong antimicrobial effects against broad-spectrum of microorganisms and *in vitro* antioxidant ability on DPPH. Our results give a perspective that *Tephromela atra* extracts and the isolated metabolites can be effectively used as a source of antimicrobial and antioxidant agents in the field of pharmaceutical industries. The bioactive metabolites from *Tephromela atra* can be alternative remedies to synthetic drugs in the pharmaceutical industry and consider as promising therapeutic agents from natural sources. By expanding this type of work, it will be possible to determine the biological activities of lichen species-metabolites and investigate their possible uses in medicine, pharmacology, and industrial areas.

## ACKNOWLEDGEMENT

The authors thank Eskisehir Technical University, Scientific Research Projects Commission for funding the isolation of lichen acids and biological studies. The authors thank all faculty and staff members of Anadolu University, Faculty of Pharmacy, DOPNALAB for High-Resolution Mass Spectroscopy (HRMS) analysis.

## Authors’ Contributions

Conceptualization, A. A., and M. Y.; Methodology, A. A., and M. Y.; Validation, M. Y., I. A., and M. C.; Investigation, A. A., and M. Y.; Writing – Original Draft Preparation, A. A.; Writing – Review & Editing, A. A., M. Y., I. A. and M. C.; Supervision, M. Y.; Funding Acquisition, M. Y. All authors read and approved the final manuscript.

## Ethics Declarations

### Consent for publication

All authors confirmed the publication of the manuscript.

### Ethical approval

Our research does not need ethical approval.

### Human and animal rights

This article does not contain any studies on human or animal subjects.

