## Supplementary Information for "Investigation of Antimicrobial and Antioxidant Activity of *Tephromela atra* Lichen and its Chemical Isolates, α-Alectoronic Acid and α-Collatolic Acid"

**for**

**Contents	page**

Copies of HRMS [ESI(+)-TOF] of for

α-alectoronic acid ………………………………………...……………………………………………………………………………............. 2

α-collatolic acid ………………………………………………………………………………………………………………………………... 3

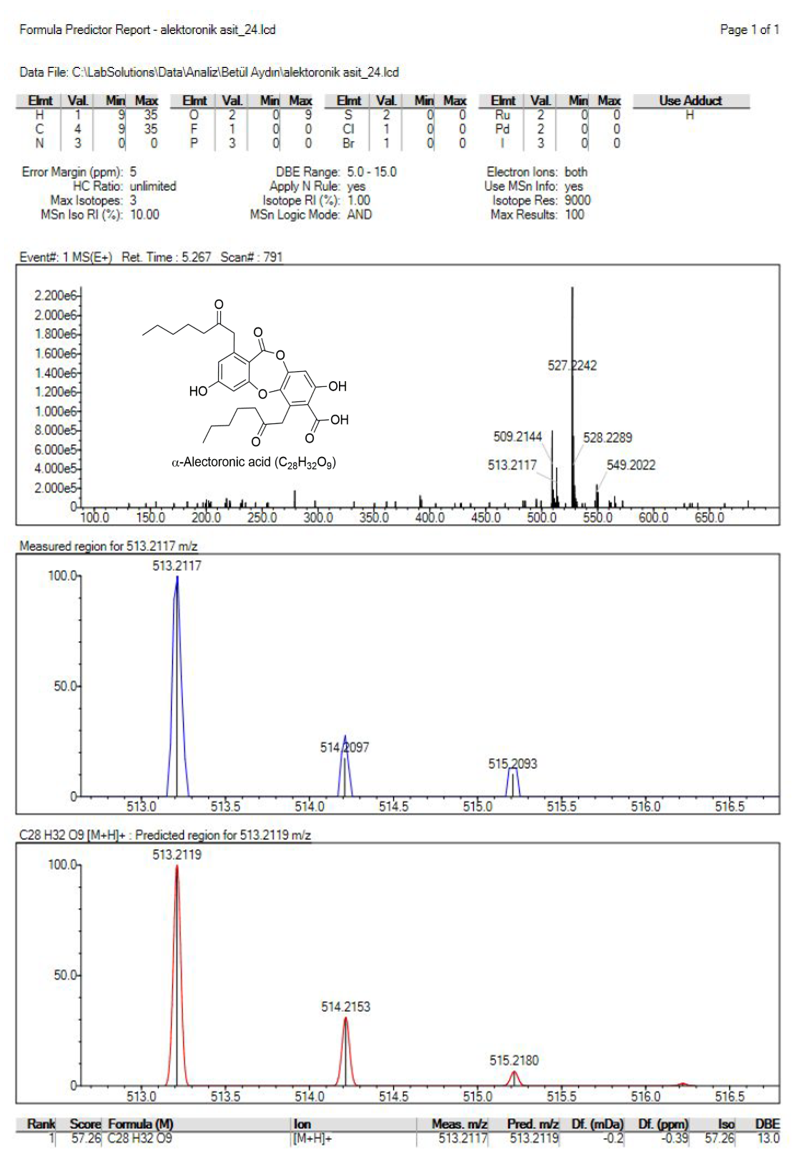

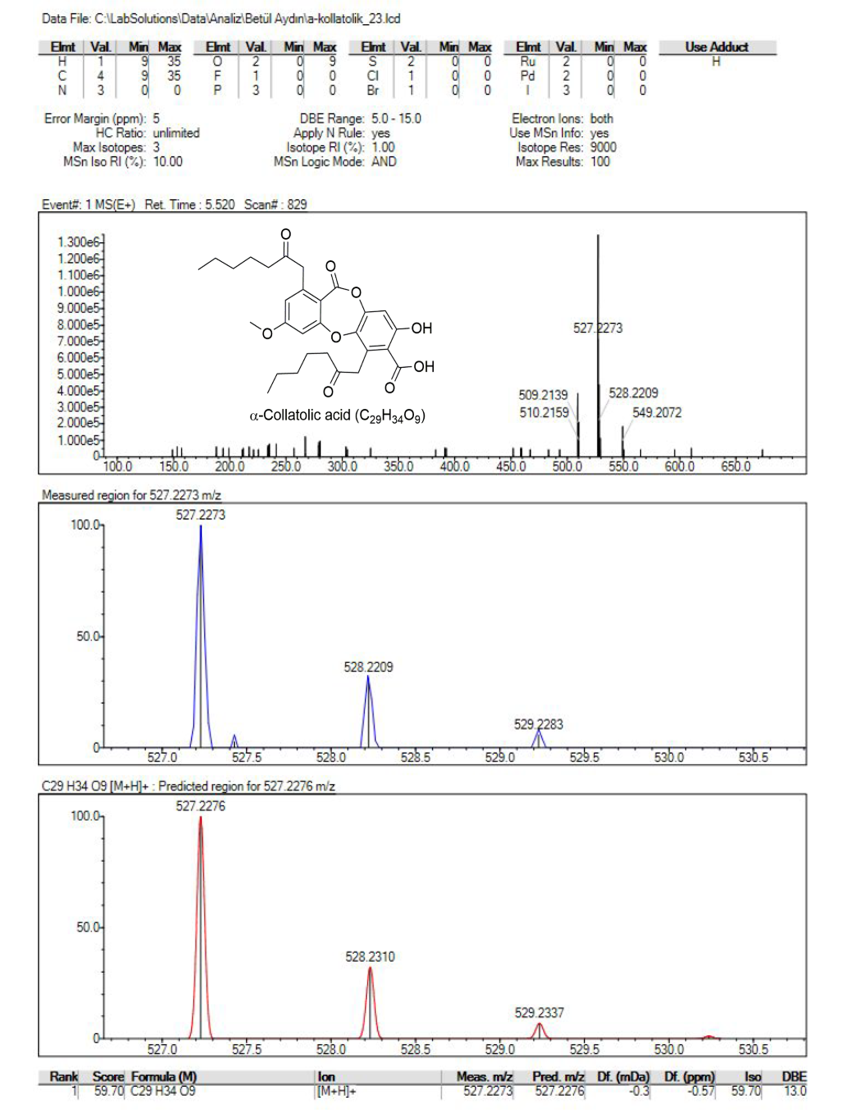
